# Infrared imaging supports warming effects of anthocyanin pigments in flowers of diverse plant taxa

**DOI:** 10.64898/2026.08.07.743445

**Authors:** Nicole M. Hughes, Joshua W. Campbell, Elizabeth D. Ragan, Sarah Jessica Forte, Natalie M. West

## Abstract

Flower color has primarily been studied in the context of pollinator attraction, although effects on thermal energy balance are also important, especially in the context of global climate change. We used infrared imaging to compare petal temperatures of white versus pigmented cultivars of ten angiosperm taxa under controlled environmental conditions. Excised sets of flowers (n= 6 sets per species) exhibiting white, light, and/or dark anthocyanin (red to purple) coloration were mounted perpendicularly to the sun at mid-day, under clear sky, low wind (<1 m s^-1^) conditions. Sunlight was filtered through either UV-transparent or UV-opaque film, and petal temperatures were measured using an infrared camera after one minute equilibration. In all species, pigmented flowers were significantly warmer than lighter-colored conspecifics. Mean differences averaged +5.3°C for darker-colored versus white morphs, +2.9°C for lighter-colored versus white. Most warming was associated with visible wavelengths, but additional warming under UV-inclusion was also observed in some species. *In situ* observations of intact landscape plants under low-wind, high-light conditions corroborated experimental results, with differences exceeding 10°C observed in some taxa. Temperature differences >7°C were also recorded for purple versus white sections of the same flower in multicolored *Viola* and *Petunia* cultivars. Follow-up experiments using dark-pink and white varieties of *Impatiens x hybrida* corroborated well-known effects of sunlight intensity and wind speed on floral temperatures, helping to explain inconsistent reports in the literature. Our results clearly demonstrate that anthocyanin pigments can have significant and dramatic impacts on floral temperatures, which could be an important factor driving evolution of flower color. In the context of climate change, floral pigments could amplify the effects of rising global temperatures, negatively impacting plant reproduction and crop yields, especially on the warmer end of species’ ranges. Changes in flower color could also potentially induce shifts in pollinator communities, which could have community-scale effects.

## Introduction

Flower coloration is one of the most conspicuous attributes of angiosperms, and the forces driving diversification of flower color are of great interest to ecologists and evolutionary biologists, as well as the horticulture industry. Anthocyanin pigments are most commonly responsible for pink, red, purple, and blue colors observed in flowers, although other pigments (e.g., carotenoids, betalains) can impart similar hues in certain taxa (reviewed in [1]). Anthocyanin-based flower color polymorphisms are more common in plants compared to polymorphisms of other floral pigments [2], which may be due to the relatively rapid rate of evolution of anthocyanin pathway genes [3].

Flower color has traditionally been thought to be selected for primarily by pollinators [4, 5, 6], and indeed, flower shape and color are often associated with certain types of pollinators (i.e., pollinator syndromes) [7]. Birds are often associated with visiting red flowers [8], whereas bees and many other insects appear less attracted to red flowers, possibly due to their lack of red photoreceptors [9, 10, 11, 12]. Changes in pollinator communities have also been shown to drive intraspecific frequencies of flower color [13]. Conversely, mutations resulting in changes in flower color also induce changes in pollinator identity [14].

Although there is much support for the idea that floral pigments play an important role in attracting pollinators, more recent studies suggest that factors influencing evolution of flower color may be more complex [15, 16, 17]. For example, most pollinating insects are generalists [18, 19], and many flowers that appear red to humans are now known to be visited by bees [20]. Flower color is also now recognized as playing a role in herbivore defense [21, 22], camouflage [23], UV protection [24], protection from fungal pathogens [21] and passive solar heating [25, 26]. This latter process, which is distinct from metabolically-driven thermogenesis (e.g., [27, 28]) is the focus of the current study.

Factors which influence floral temperatures are important to understand because they influence several key aspects of plant reproduction, physiology, and ecology (Fig. 1). This is especially important in the context of global climate change, where rapidly increasing global air temperatures have been linked with more extreme heatwaves and drought events [29]. Examples of processes known to be temperature-sensitive include flower opening, vaporization of pollinator-attracting volatiles, pollen development and germination, pollen tube formation, fertilization, and ovule development [30, 31, 32, 33, 34, 35, 36]. “Heat rewards” are also known to significantly influence pollinator visitation, especially in colder climates [37, 38, 39]. Heat rewards can raise an insect’s metabolic rate and potentially increase foraging activity when ambient temperatures are low [40, 41]. Bumble bees were shown to prefer to feed on warm sugar solutions compared to room temperature sugar solutions and would use flower color to differentiate which artificial flowers were warmer and contained the warmer sugar solution [39]. Similarly, cooler flowers provide pollinators an escape from intense heat in warmer climates, and some bee species have been shown to prefer cooler nectar when ambient temperatures exceed 34°C [42]. Heat rewards would not be expected to play a direct role in vertebrate (e.g., birds, bats) visitation. However, nectar secretion can be reduced at low temperatures and floral warming may increase the production of nectar [43] and diffusion of volatiles [44]. Thus, although the vertebrate pollinator is not visiting for the heat reward, floral warming may indirectly lead to an increase in vertebrate flower visitation.

**Figure 1.**
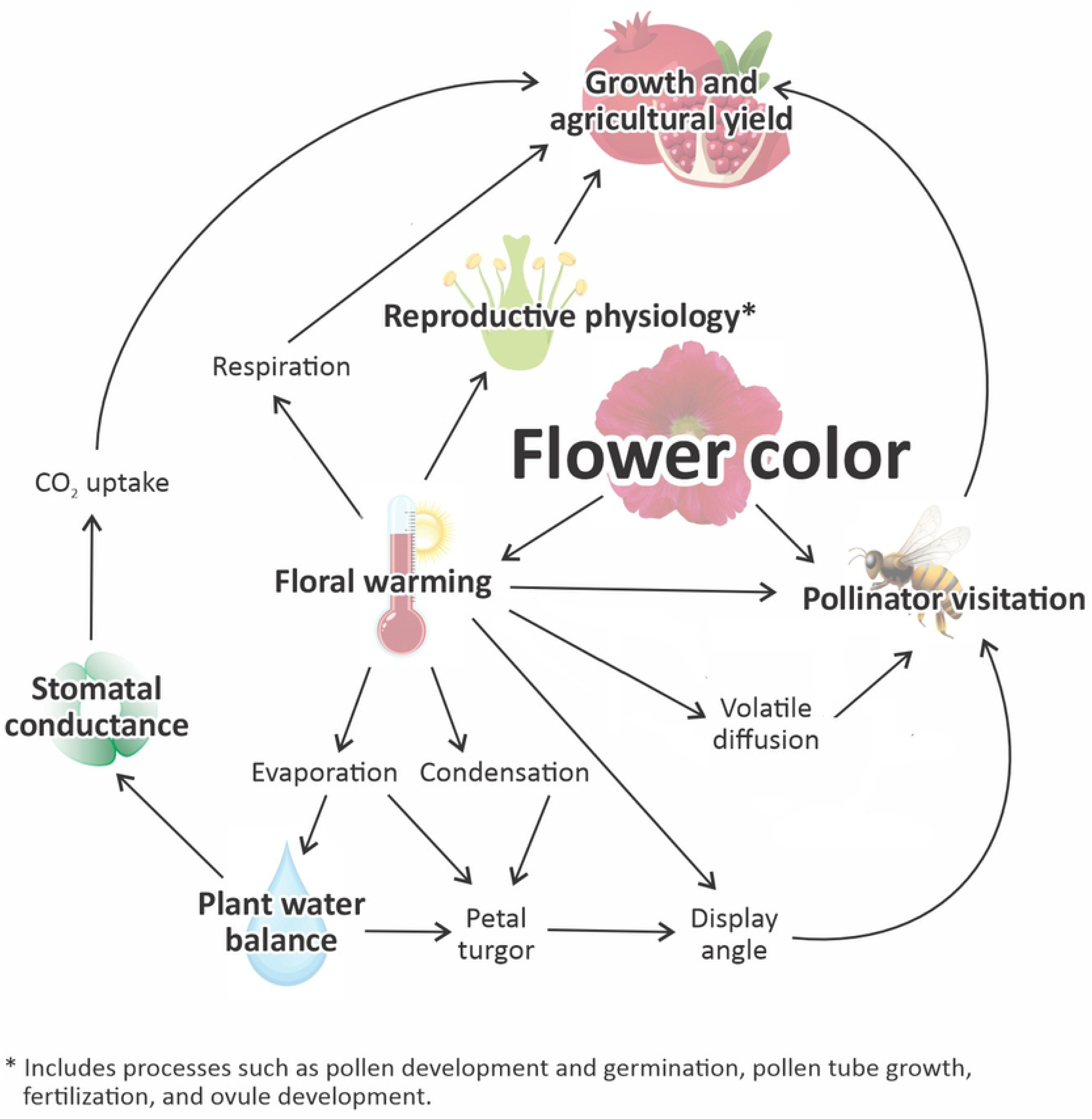
Flow chart illustrating the range of physiological and ecological factors influenced by flower color.

Like other polyphenols, anthocyanins absorb strongly in the ultraviolet wavebands [45], but are unique in their additional absorbance of visible light, primarily in the green wavebands (but see [46]). The addition of various chemical moieties can further enhance molar absorptivity of anthocyanins. For example, acylated anthocyanins produced in drought-stressed leaves of *Arabidopsis thaliana* absorbed 1.8 times more visible light and nearly 3 times more UV-B than equivalent concentrations of the most common anthocyanin—cyanidin-3-glucoside [47]. Although the visible spectrum has traditionally been understood as the only waveband relevant to plant thermal energy balance [48], recent studies have demonstrated measurable and significant contributions of ultraviolet UV wavelengths as well [26].

While thermal effects of pigments on plant tissues may seem obvious, the literature paints a more conflicting picture ([41] and citations therein). In support of a warming effect, multiple studies have reported darkly-pigmented flowers to be between 2-6 °C warmer than white conspecifics [26, 49, 50, 51, 52]. Colorful flower morphs of some species are also more common in cooler climates or at higher elevations [50, 51, 53, 54]. Darkly-pigmented flowers in plant species which are phenotypically plastic for flower color have been reported to occur at higher frequencies than lighter flowers during cooler periods and vice versa [25, 55]. In flowers that close petals around young developing fruits, fruits tend to be warmer when exposed (abaxial) petal surfaces are purple rather than white [56].

Yet, results to the contrary have also been reported. White flowers of some species polymorphic for flower color have been reported at higher frequencies under *cooler* conditions or *higher* elevations than darker color morphs [57, 58, 59]. Several authors have also reported no significant difference in petal temperatures of pigmented versus white flowers [58, 60, 61, 62]. It is possible, however, that the lack of temperature differences observed in these latter studies could be the product of biophysical confounds in experimental design. For example, most studies that did not observe a temperature difference also did not report controlling for environmental parameters known to impact plant temperatures, such as wind, sunlight intensity, and/or compass direction, e.g., [60, 61, 62]. Others did not directly measure the temperature of the flower itself, but rather the temperature of air spaces within the flower [30, 63]. It is well known that thermal conductivity is lower in air compared to water-based tissues. Still others tested for warming effects using extremely low light levels, insufficient to induce significant warming. In [30], the light source used to heat flowers consisted of two desk lamps with a combined output of only 50 µmol m^-2^ s^-1^ photosynthetic photon flux density (PPFD) (compare this to PPFDs near 1500-2000 µmol m^-2^ s^-1^ reported at solar noon in temperate and tropical latitudes, e.g., [64, 65]. Temperature differences between white and pigmented flowers are also greatest under increased irradiance [51, this study], which could account for a lack of difference in this latter example. Finally, Shrestha *et al*., [61] compared flower temperatures between unrelated taxa, which introduces biophysical confounds related to differences in flower morphology [41]. The only study that we are aware of which reports no significant effect of flower color on temperature despite using within-species comparisons and controlling for sunlight intensity and wind was Mu *et al*. [58]. In this case, white flowers of *Gentiana leucomelaena* did not differ in temperature from blue-colored morphs under clear and non-windy, windy, or overcast conditions. However, the lack of temperature difference in this case may be attributed to the extremely small size of the flower (ca. 1 cm in diameter). It is known that smaller surfaces return to air temperature more rapidly due to greater wind flow over the surfaces, which promotes a more turbulent boundary layer and increased convective heat loss [48].

The objective of the current study was to use infrared imaging to test the hypothesis that anthocyanin pigments increase petal temperatures under bright, low-wind conditions. To control for interspecific variation in flower morphology, ten flowering plant species were selected based on the availability of conspecific cultivars with variable amounts of floral anthocyanins. To control for environmental factors which also influence flower temperature, experiments were conducted under clear skies at mid-day (1100-1300 h), under low-wind (< 1 m s^-1^) conditions. Our analysis also took into account morphological factors, including petal thickness and corolla diameter. To differentiate visible from UV warming effects, measurements were made under UV-opaque vs. UV-transparent films. Because controlled environments do not necessarily translate into real world conditions, flower temperatures of intact plants under natural conditions were also measured to corroborate experimental results.

## Materials and Methods

### Plant material

Plant material used in all experiments was obtained from the Mariana H. Qubein Arboretum and Botanical Gardens on the campus of High Point University, North Carolina USA (35° 59’ 6.0518’’ N, 80° 0’ 6.6679’’ W). Species were selected based on the availability of two or more cultivars differing in floral anthocyanin content (low-high). In some species, different color morphs were produced on the same individual plant (e.g., *Rhododendron* ‘Conlep’ Autumn Twist, *Salvia microphylla*). Species used for experiments are illustrated in Fig. 2, and included azaleas-*Rhododendron* ‘Conlep’ Autumn Twist (light pink and dark pink flowers), and ‘Robleg’ Autumn Angel (white); dark pink, light pink, and white Madagascar periwinkle-*Catharanthus roseus,* red, pink, and white wax begonias-*Begonia x semperflorens-cultorum;* dark pink, light pink, and white impatiens-*Impatiens x hybrida* SunPatiens®; red and white baby sage-*Salvia microphylla*; light purple and white rose of Sharon-*Hibiscus syriacus;* dark purple, light purple, and white pansies-*Viola x wittrockiana;* dark pink, and light purple, and white crepe myrtle-*Lagerstroemia indica*; red, pink, and white star flower-*Pentas lanceolata;* dark purple, light purple, and white Angelonia-*Angelonia angustifolia*.

**Figure 2.**
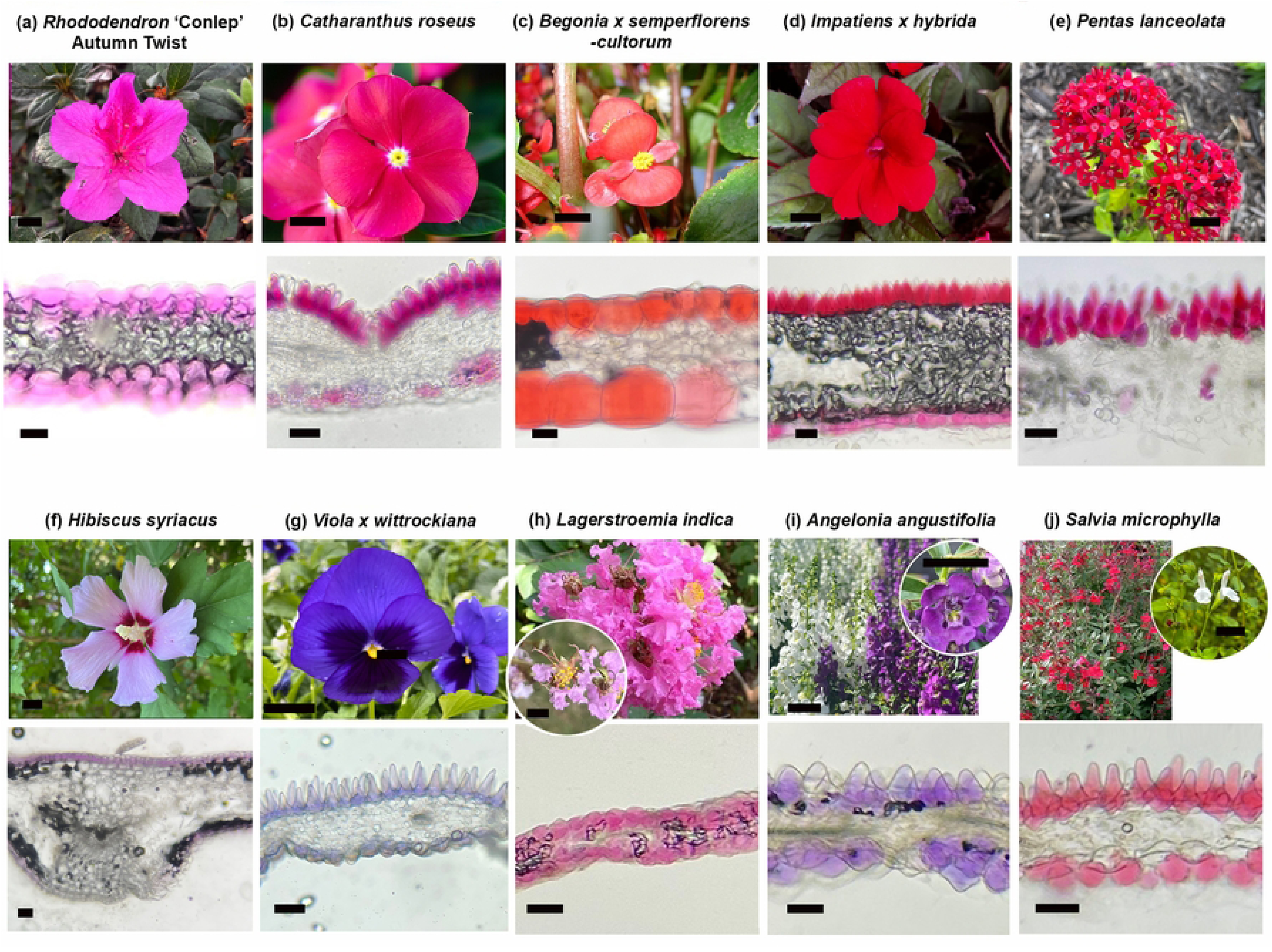
Species used in flower temperature experiments with their petal cross sections. Pictured are azalea, *Rhododendron* ‘Conlep’ Autumn Twist (a), Madagascar periwinkle, *Catharanthus roseus* (b), wax begonia, *Begonia x semperflorens-cultorum* (c), impatiens, *Impatiens x hybrida* SunPatiens® (d), star flower, *Pentas lanceolata* (e), rose of Sharon, *Hibiscus syriacus* (f), pansy, *Viola x wittrockiana* (g), crepe myrtle (individual floret), *Lagerstroemia indica* (h), Angelonia (individual floret), *Angelonia angustifolia* (i), and baby sage, *Salvia microphylla* (j). Scalebar for photos is 1 cm, and 200 µm for microscopic cross sections.

Diameter of corollas used in temperature experiments were measured for all experimental plants using imageJ software (NIH, USA). For measurements of petal thickness, 8-12 random flowers of each species were placed in a closed container with a wet paper towel to rehydrate overnight, to maximize turgor pressure. The next day, petal thickness was measured between two glass coverslips using digital calipers using individual petals (for small flowers) or a 1 cm^2^ piece taken from the center of the petal (for larger flowers).

For observations of intact plants in the field, measurements were taken on co-occurring color morphs on the High Point University campus on clear, low wind days. Air temperature was measured with a Kestrel 2500 weather meter, and photosynthetic photon flux density (PPFD) was measured using an LI-250A light meter (Li-COR, Inc., Lincoln, NE, USA).

### Microscopy

For microscopic imaging, fresh flowers were excised in the morning, and kept in a cool, moist, air-tight container prior to analysis later that day. Petals were sectioned using an Oxford vibratome by placing tissues within a cube of fresh potato and viewed under light microscopy. Images were captured with an iPhone 5 through the ocular lens (Apple, Cupertino, CA).

### Anthocyanin content

To estimate anthocyanin concentration in flower petals, three hole punches (32.4 mm^2^ area each), or in the case of very small flowers, three flowers, were excised from each plant and freeze-dried. Dried tissue was massed and extracted in 2 mL methanol:acetic acid:water (70:23:7 by vol) for 48 h at 4°C in the dark. Anthocyanin concentration was measured spectrophotometrically as absorbance at 530 nm and normalized by dry mass.

### Infrared imaging experiments

Flowers were removed from plants and kept in an air-tight container containing a wet paper towel until measurement within 3-5 h of excision. Measurements were taken only on clear or mostly clear (<5% cloud cover) days to minimize the risk of diffuse photons striking flowers without first passing through UV inclusion/exclusion films. Wind was minimized by conducting experiments in an outdoor, 3.3 m tall brick-walled enclosure that was open to the sky, on low-wind days (ambient wind speeds < 2 m s^-1^). For each measurement, a set of conspecific flowers with low, medium, and/or high anthocyanin content were mounted perpendicularly to the sun, in random order, into a row of holes in a white cardboard piece (30.5 cm x 24 cm) oriented perpendicularly to the sun. For species with many small florets arranged in inflorescences, individual florets were measured. The cardboard piece was placed behind a UV-inclusion (Aclar) or UV-exclusion (Courtguard) film, which was also angled perpendicularly to the sun. Flowers were shaded during set-up. After exactly 1 min exposure to full ambient sunlight (determined prior to experimentation as sufficient to reach steady-state warming; data not shown), thermal images of the set of flowers were taken using a Fluke TiR110 thermal imaging camera (Fluke Corp., Everett, Washington, USA). Following measurement, flowers were shaded until they had cooled back down to air temperature, and then measured again under the alternate UV treatment (typically within <3 min). Because water loss could potentially impact temperature via changes in turgor pressure and/or stomatal conductance, the order of UV inclusion/exclusion treatments alternated between replicates. Temperature in three standardized points on each flower petal were averaged using Fluke SmartView® imaging software. Additionally, temperatures of central reproductive structures was estimated for larger species by averaging points along a line through the center of the flower using SmartView software. Interior temperatures were included for *Begonia, Hibiscus, Impatiens,* and *Catharanthus* spp.

### Sunlight and wind effects

Five pairs of dark pink and white *Impatiens x hybrida* SunPatiens® of similar size were utilized for testing the effects of sunlight intensity and wind on flower temperatures. Flowers were excised in the morning and stored in an air-tight plastic container with a wet paper towel until measurement within 3-5 h. To control for wind and sunlight, measurements took place indoors in front of a south-facing window on a sunny day. These experiments were conducted in July at mid-day (i.e., three hours before and after solar noon). Petioles of excised flowers were held in place using miniature alligator clamps attached to a weighted white resin cube (7.5 cm^3^), so that flowers could be oriented perpendicularly to the sun. To protect tender petioles from being broken by the sharp teeth of the miniature alligator clamps, several layers of lab tape were wrapped around each jaw.

To quantify the effect of sunlight intensity on flower temperature, infrared images were taken under varying degrees of shade provided by neutral density shade screens placed between pairs of pink and white flowers and the window. Flowers were held in place using invitation holders, consisting of a block with a vertical wire and terminal alligator clip. Sunlight intensity was measured using a LI-250A light meter. Infrared images were captured after 1 min at each light level.

For wind experiments, a small desk fan with three levels of intensity was set on the end of 2.25 m long paper “runway” facing a window, which transmitted 1000-1100 µmol m^-2^ s^-1^ PPFD at the time of the experiment. Points along the center of the runway were marked at intervals that corresponded with different wind speeds as measured using a Kestrel 3500 handheld weather meter. Dark pink and white flowers were paired by size, held in place using the alligator clamps as described in the previous section, and placed along each point in the runway until a steady state temperature was reached at each fan speed, at which time a measurement was taken with the Fluke TiR110 thermal imaging camera.

### In situ measurements

During May and June 2026, infrared images of intact plants *in situ* displaying different colored flowers and occurring side-by-side were captured using the Fluke TiR110. Prior to each measurement, sunlight intensity was measured with a LI-250A light meter, while wind speed and air temperature were measured using a Kestrel 3500 handheld weather meter. Because quality of visible images captured by the camera were low resolution, an iPhone X camera was used to take photos of the plants from the same angle and distance as infrared images.

### Statistics

To evaluate the influence of different factors on temperature response, we used a linear mixed effects model (R package lme4, ‘lmer’) with Type II SS (R package car, ‘Anova’). Individuals nested in species were included as random blocks, and each species had one white (control), one “light” (intermediate anthocyanin value) morph, and/or one “dark” (higher anthocyanin value) morph. We compared fitted estimates to assess the relative influence of each individual term; in the case of color, we compared among colors, and with white. We examined single-term deletions (‘drop1’ in R base package) to compare the contribution of each term to model fit and excised terms that did not result in more than a two-point decrease in AIC compared to an intersect-only model. We included color as both a categorical variable and as a quantitative variable, the latter using mean absorption at 530 nm. Pairwise differences among colors were further validated with post-hoc Tukey tests (R package multcomp, ‘glht’). Analyses were performed in R v3.6.3 (R Core Team 2020).

Within-species comparisons of petal temperatures between different colored morphs were analyzed using ANOVA with Tukey’s HSD *post hoc* test, with individuals blocked by set (one set = one white, one lightly-pigmented, and one darkly-pigmented conspecific cultivars of similar size tested at the same time). Ultraviolet inclusion and exclusion were tested as separate ANOVAs. Within each color treatment of each species, a paired t-test was used to compare petal temperatures under UV-inclusion versus exclusion. For the four species where interior temperatures could be additionally measured, a MANOVA with Tukey’s HSD *post hoc* test was used to measure the effect of color on petal and interior temperatures. A paired t-test was also used to compare petal and interior temperatures of each color morph. Significance was determined as p<0.05. Due to biological and genetic differences among species we also report marginal significance up to p<0.1, to recognize potentially biological significantly patterns.

## Results

### Microscopy

Microscopic examination of flower petals (Fig. 2) showed that anthocyanins were concentrated in both the adaxial (upper) and abaxial (lower) epidermis of petals equally in six species (*Rhododendron* sp.*, Salvia microphylla, Viola x wittrockiana, Hibiscus syriacus, Angelonia angustifolia,* and *Lagerstroemia indica*), and primarily to exclusively adaxially located in four species (*Begonia x semperflorens-cultorum, Impatiens x hybrida* SunPatiens ®*, Pentas lanceolata, Catharanthus roseus*). Dermal papillae were clearly visible under 100x in *A. angustifolia, Impatiens x hybrida* SunPatiens®*, P. lanceolata, C. roseus, S. microphylla,* and *Viola x wittrockiana.* Mesophyll cells within the petals were mostly clear, but low levels of anthocyanins were occasionally observed in cell layers adjacent to the epidermis (*Begonia x semperflorens-cultorum, Impatiens x hybrida* SunPatiens ®, and *L. indica)*.

### Anthocyanin content

In all ten plant species examined, visually darker (e.g., dark red/purple) colored flowers had significantly higher anthocyanin content compared to those which were visually lighter (pink or light purple); white flower morphs contained zero or only trace amounts of anthocyanins, and these levels were always significantly lower than those in pigmented cultivars (Table 1).

**Table 1.** List of plant taxa, flower petal color (as seen by the human eye), mean petal thickness, corolla diameter, and anthocyanin content (with SE) for different color morphs used for controlled temperature experiments (n=6-10). Mean warming effects relative to white conspecifics are included in the presence (+) and absence (-) of UV light (Δ Temp) for light and dark color morphs. P-values are from one-tailed, paired t-test comparing petal temperatures under UV+ versus UV-for white, light, and dark color morphs. Significant differences are highlighted in bold.

| Species<br>(Family) | Petal<br>color | Petal<br>thickness<br>(mm) | Corolla<br>diameter<br>(cm) | Anthocyanin<br>$A^{530}$/g | $\Delta$ Temp<br>(°C)<br>UV+ | $\Delta$ Temp<br>(°C)<br>UV- | UV<br>effect<br>(p-value) |
| --- | --- | --- | --- | --- | --- | --- | --- |
| *Rhododendron*<br>sp.<br>(Ericaceae) | White | 0.22 | 7.7 | 0.095 (0.1) | — | — | **\*0.048** |
|  | Light | (0.05) | (0.1) | 6.7 (2) | +2.7 (0.9) | +3.3 (0.4) | 0.30 |
|  | Dark |  |  | 39 (0.5) | +6.0 (0.4) | +5.2 (0.7) | 0.23 |
| *Catharanthus*<br>*roseus*<br>(Apocynaceae) | White | 0.21 | 4.4 | 0.18 (0.07) | — | — | 0.44 |
|  | Light | (0.01) | (0.04) | 9.3 (6) | +3.3 (0.6) | +2.9 (0.3) | 0.23 |
|  | Dark |  |  | 86 (20) | +7.9 (0.8) | +8.2 (0.6) | 0.46 |
| *Begonia x semper-*<br>*florens-cultorum*<br>(Begoniaceae) | White | 0.33 | 2.4 | 0.0 (0) | — | — | 0.084 |
|  | Light | (0.02) | (0.04) | 100 (46) | +2.5 (0.3) | +1.9 (0.3) | **\*0.039** |
|  | Dark |  |  | 1200 (170) | +4.7 (0.4) | +4.0 (0.3) | 0.070 |
| *Impatiens x hybrid*<br>SunPatens®<br>(Balsaminaceae) | White | 0.48 | 5.5 | 0.0 (0) | — | — | 0.17 |
|  | Light | (0.02) | (0.1) | 20 (7) | +4.4 (0.05) | +3.5 (0.4) | 0.077 |
|  | Dark |  |  | 125 (20) | +7.9 (0.8) | +8.2 (0.6) | 0.13 |
| *Pentas lanceolata*<br>(Rubiaceae) | White | 0.25 | 1.3 | 0.32 (0.3) | — | — | 0.093 |
|  | Light | (0.01) | (0.03) | 48 (5) | +1.9 (0.4) | +2.5 (0.4) | 0.11 |
|  | Dark |  |  | 193 (23) | +3.4 (0.2) | +3.5 (0.2) | 0.08 |
| *Hibiscus syriacus*<br>(Malvaceae) | White | 0.30 | 7.7 | 0.0 (0) | — | — | 0.23 |
|  | Light | (0.01) | (0.2) | 13 (6) | +3.1 (0.4) | +3.6 (0.3) | 0.47 |
| *Viola*<br>*x wittrockiana*<br>(Violaceae) | White | 0.21 | 3.1 | 0.70 (0.3) | — | — | 0.12 |
|  | Light | (0.01) | (0.1) | 85 (16) | **+5.0 (1)** | **+4.2 (0.7)** | **\*0.009** |
|  | Dark |  |  | 377 (25) | +7.6 (0.7) | +7.6 (0.8) | 0.16 |
| *Lagerstroemia*<br>*indica*<br>(Lythraceae) | White | 0.077 | 3.3 | 0.56 (0.4) | — | — | 0.26 |
|  | Light | (0.01) | (0.2) | - | +2.2 (0.5) | +2.3 (0.8) | **\*0.017** |
|  | Dark |  |  | 110 (20) | +3.1 (0.6) | +2.8 (0.5) | 0.061 |
| *Angelonia*<br>*angustifolia*<br>(Plantaginaceae) | White | 0.14 | 1.0 | 1.5 (0.1) | — | — | 0.28 |
|  | Light | (0.01) | (0.05) | 15 (2) | +1.6 (0.5) | +1.4 (0.2) | 0.36 |
|  | Dark |  |  | 37 (5) | +2.0 (0.4) | +1.3 (0.3) | 0.45 |
| *Salvia nemorosa*<br>(Lamiaceae) | White | 0.16 | 1.3 | 0.054 (0.05) | — | — | 0.055 |
|  | Red | (0.01) | (0.04) | 27 (3) | +2.2 (0.5) | +2.3 (0.8) | 0.10 |
| **Mean (SE):** |  |  |  | (lighter color) | **+2.9 (0.3)** | **+2.8 (0.3)** |  |
| **Mean (SE):** |  |  |  | (darker color) | **+5.3 (0.8)** | **+5.1 (0.9)** |  |

### Effects of flower color on temperature

The effect of corolla color on petal temperature was tested as both a categorical variable and as a quantitative variable using mean anthocyanin content (A530 g^-1^ dry tissue), but there were no qualitative differences in model results (data not shown). Petal color and UV treatment significantly influenced temperature in the mixed effects models (color: p < 0.001, χ^2^ = 215.42, 5df; UV: p = 0.004, χ^2^ = 12.59, 1 df), while flower diameter, dermal papillae, and petal thickness did not (diameter: p = 0.43, χ^2^ = 0.51, 1 df; papillae: p = 0.40, χ^2^ = 0.71, 1 df; thickness: p = 0.12, χ^2^ = 2.39, 1 df). These results were largely mirrored in the single term deletions. However, including dermal papillae marginally improved (reduced) the Akaike information criterion (AIC) value of the final model (data not shown).

Within-species comparisons of infrared temperatures measured under high-light, low-wind conditions showed that white petals were consistently significantly cooler than petals of conspecifics containing anthocyanins, and that darker-colored petal cultivars were always significantly warmer than lighter colored conspecifics (p < 0.001 for all pairwise comparisons with white; all regression estimates >2 units/degrees higher relative to white) (Figs. 3, 4). On average, lightly-pigmented (light pink/purple) flower petals were 2.9°C warmer than white conspecifics under UV inclusion, while more darkly-pigmented morphs (dark pink/red/purple) were 5.3°C warmer (Table 1).

**Figure 3.**
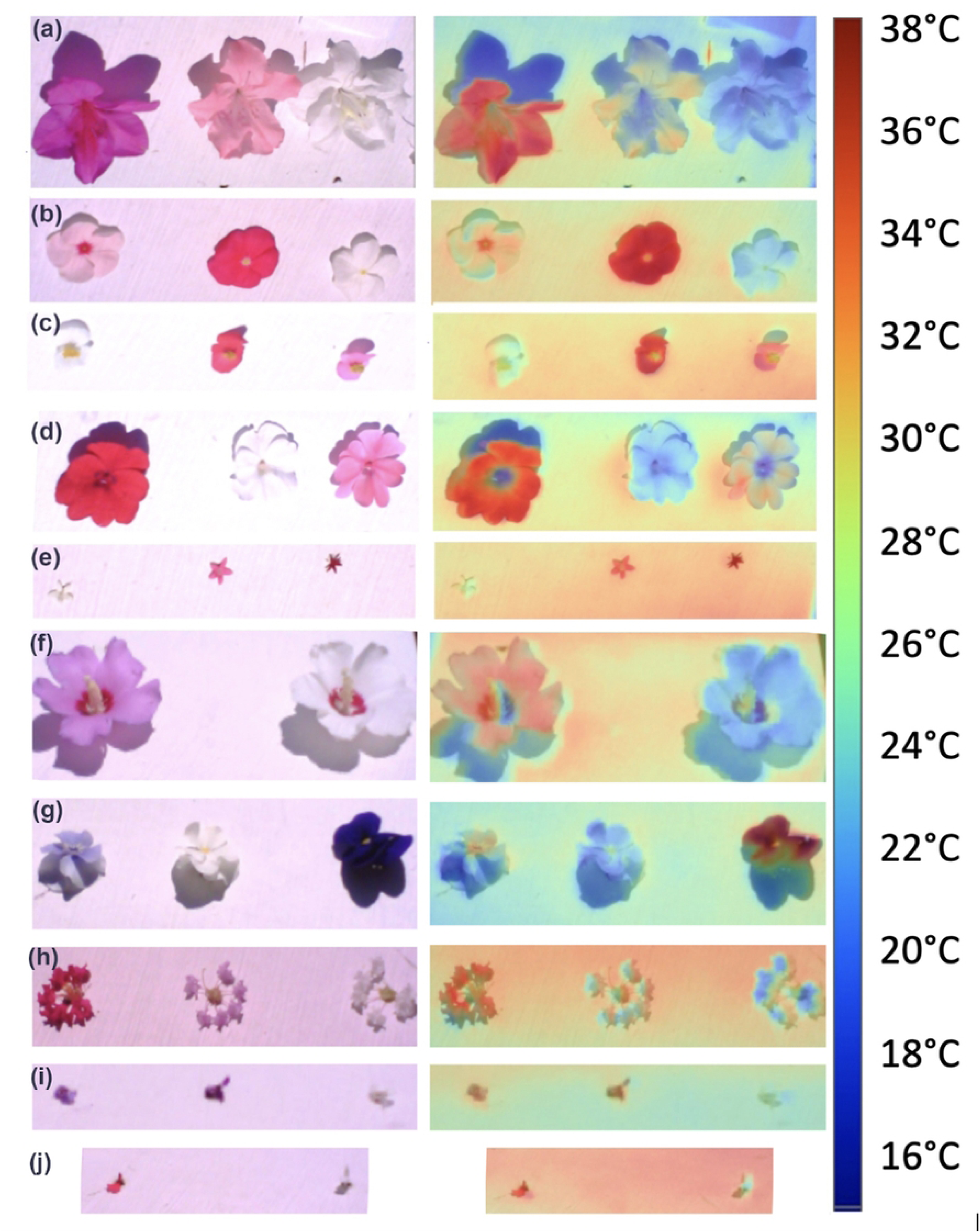
Visible (left column) and infrared (right) photos of individual replicate sets of flowers differing in anthocyanin concentration taken using Fluke TiR110 thermal imaging camera. Photos were taken following 1 min exposure to full sunlight (>1300 µmol m^-2^ s^-1^) and low wind (< 1 m s^-1^). Order of color morphs from left to right was randomized between trials. Pictured, from top to bottom, are azalea, *Rhododendron* ‘Conlep’ Autumn Twist (a), Madagascar periwinkle, *Catharanthus roseus* (b),wax begonia, *Begonia x semperflorens-cultorum* (c), Impatiens, *Impatiens x hybrida* SunPatiens*®* (d), star flower, *Pentas lanceolata* (e), rose of Sharon, *Hibiscus syriacus* (f), pansy, *Viola x wittrockiana* (g), crepe myrtle (individual floret), *Lagerstroemia indica* (h), Angelonia, *Angelonia angustifolia* (i), and baby sage, *Salvia microphylla* (j).

**Figure 4.**
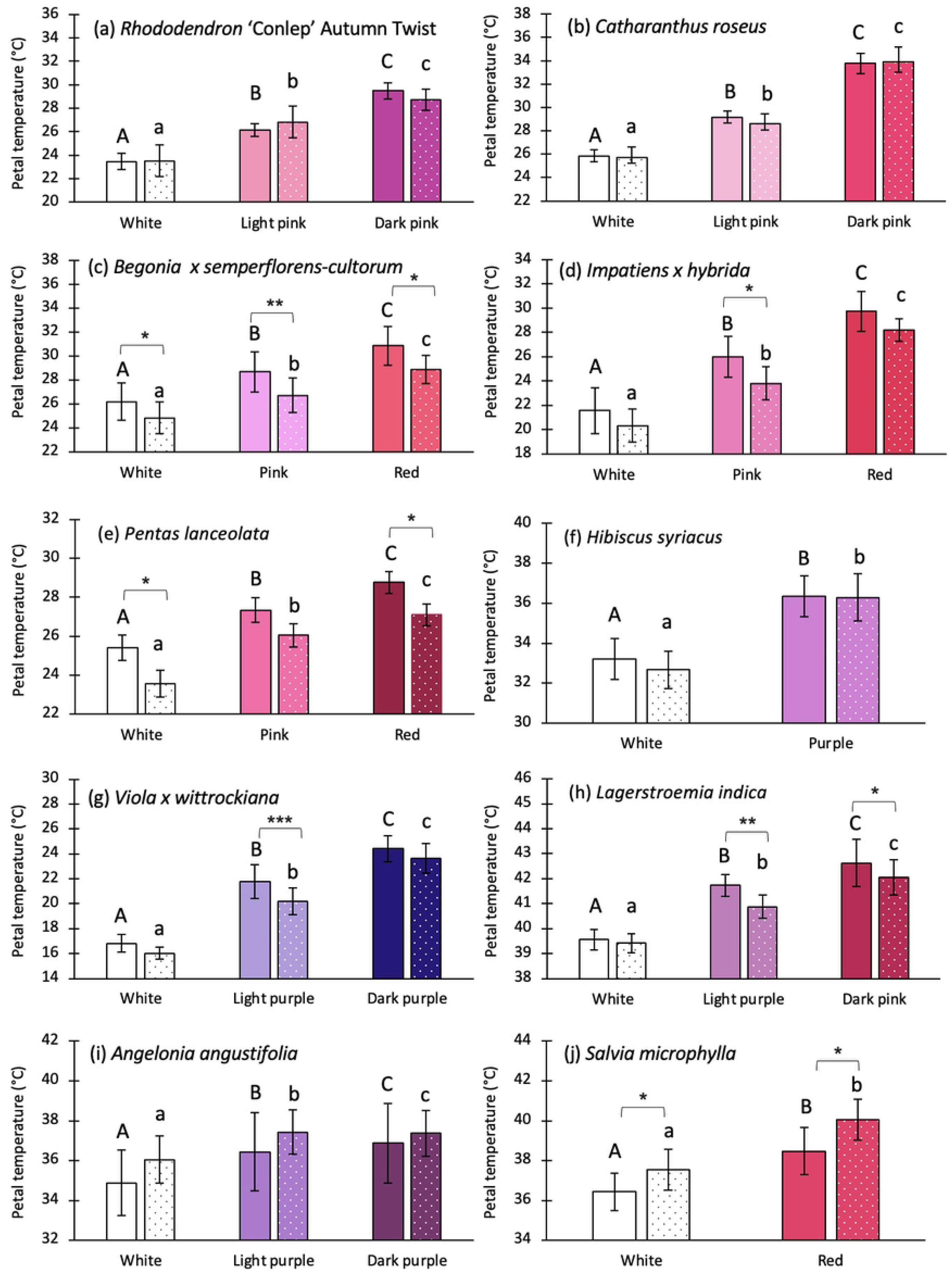
Petal temperatures from experiment depicted in Figure 3. Bars represent means of 5-6 replicates ± SE. Solid bars are petal temperatures with UV-inclusion, white-spotted bars UV-exclusion. Within-species analyses utilized ANOVA (blocked by set) to compare temperatures of the two/three color varieties. Separate analyses were run for UV-inclusion and UV-exclusion. Letters above bars represent results from Tukey’s *post hoc* test for UV-inclusion (capital letters) and UV-exclusion (lower case). Bars with different letters are significantly different at p<0.05. All *post hoc* results are the same as shown in (a). Asterisks illustrate significant (black *, p<0.05) or marginally significant (gray *, p<0.1) differences between colored flowers with and without UV as determined by within-color paired t-tests. Colors of bars match colors of flower petals tested as seen by the human eye.

Within-species paired t-tests for petal temperatures under UV inclusion versus exclusion showed significant differences only in some species. In wax begonia (*Begonia x semperflorens-cultorum*), all petal colors (including white) were significantly or marginally significantly warmer under UV inclusion (white p=0.084; pink p=0.038; red p=0.070). In impatiens *(Impatiens x hybrida*), pink flowers showed marginally significant warmer temperatures under UV inclusion (p=0.077). For pentas (*Pentas lanceolata*), both white and red florets were marginally significantly warmer under UV inclusion (p=0.093 for white, and p=0.083 for red). For pansy (*Viola x wittrockiana*), light purple flowers were significantly warmer under UV (p=0.009), but no significant differences were observed for white (p=0.12) or dark purple (p=0.16) cultivars. Individual light pink and dark red crepe myrtle (*Lagerstroemia indica*) florets were warmer under UV inclusion (p=0.017 and 0.061 respectively), while no difference was observed for white flowers (p=0.26). Finally, red and white *Salvia nemorosa* were significantly or moderately significantly warmer under UV inclusion (p=0.055 for white, p=0.010 red).

The central, interior tissues of flowers were significantly cooler than petals in nearly all lightly to darkly colored flowers, with the exception of pink *Catharanthus roseus* (Fig. 5). These results were surprising, since one might assume a thicker boundary layer to form within the throat of the flower and within the boundary layer of the larger flower. We observed that often these central tissues glistened with moisture, although we did not determine whether it was water or nectar. No significant differences in petal versus interior temperatures were observed in white cultivars of three of the four species, with the exception of *Impatiens x hybrida*. Interior temperatures were significantly warmer in pigmented compared to white conspecifics in three of the four species, and warming effects increased with increasing anthocyanin content.

**Figure 5.**
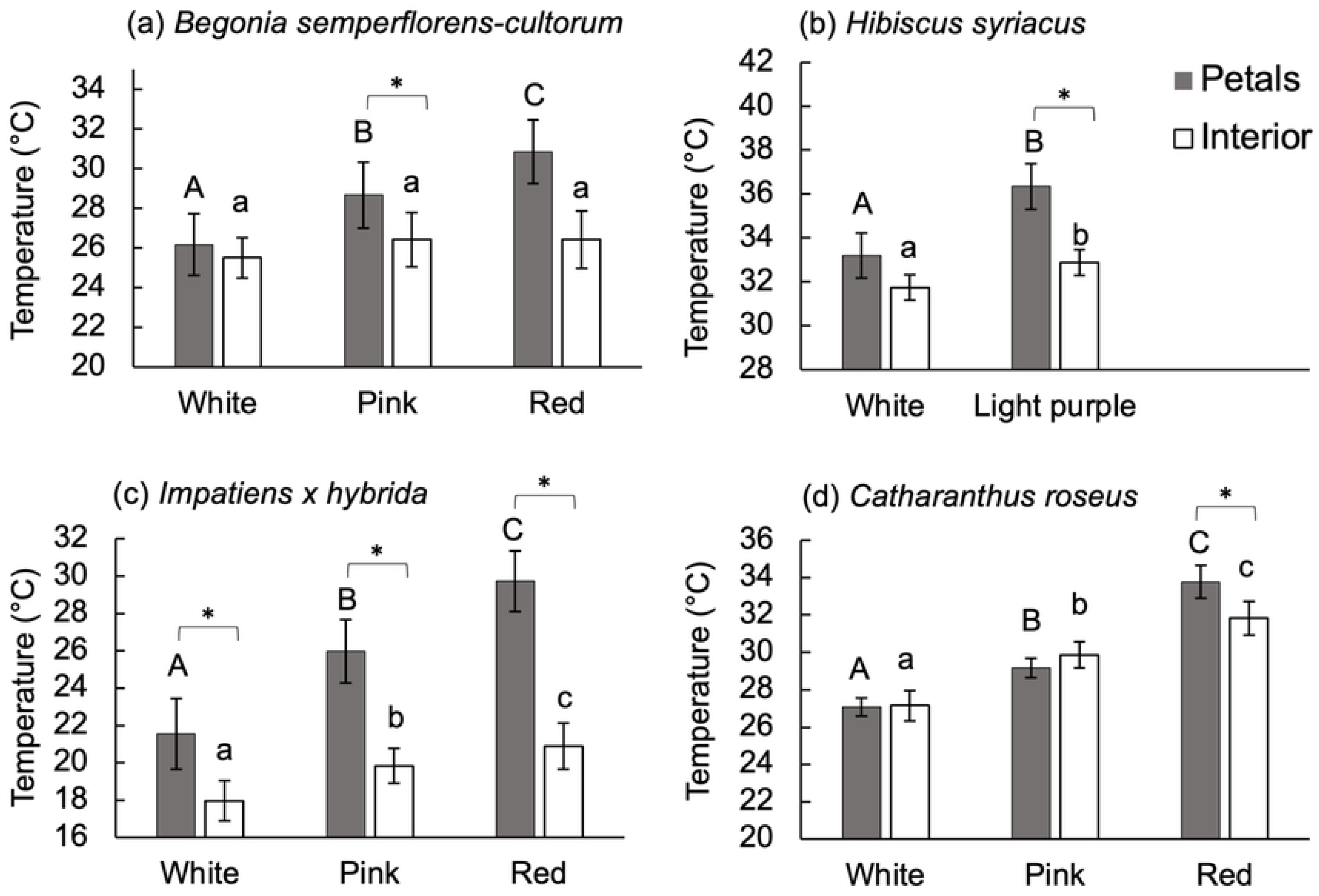
Petal versus interior tissue temperatures of select taxa. Interior tissues refer to the centers of the flowers where reproductive organs occur. Bars represent means of 5-6 replicates ±SE. Solid bars represent petal temperatures, open bars, interior tissue temperatures. Only results for UV-inclusion are shown. Within-species analyses utilized MANOVA (blocked by set) to compare temperatures of the two/three color varieties. Letters above bars represent results from Tukey’s *post hoc* for petals (capital letters) and interiors (lower case). Bars with different letters are significantly different at p<0.05. Asterisks illustrate significant (*), within-color differences between petal and interior temperatures, as determined by a one-tailed, paired t-test.

### Effects of wind and sunlight intensity on flower temperatures

Incident sunlight intensity had a significant, positive correlation with petal temperatures in white and dark pink *Impatiens x hybrida* flowers (p<0.001 for both; Fig. 6a). However, while white flowers only exhibited ca. 1°C warming between low and high irradiance, dark pink flowers warmed by 5.4°C on average. Accordingly, there was a significant, positive correlation between the difference in petal temperature between white and dark pink flowers and sunlight intensity (p<0.001; Fig. 6b).

**Figure 6.**
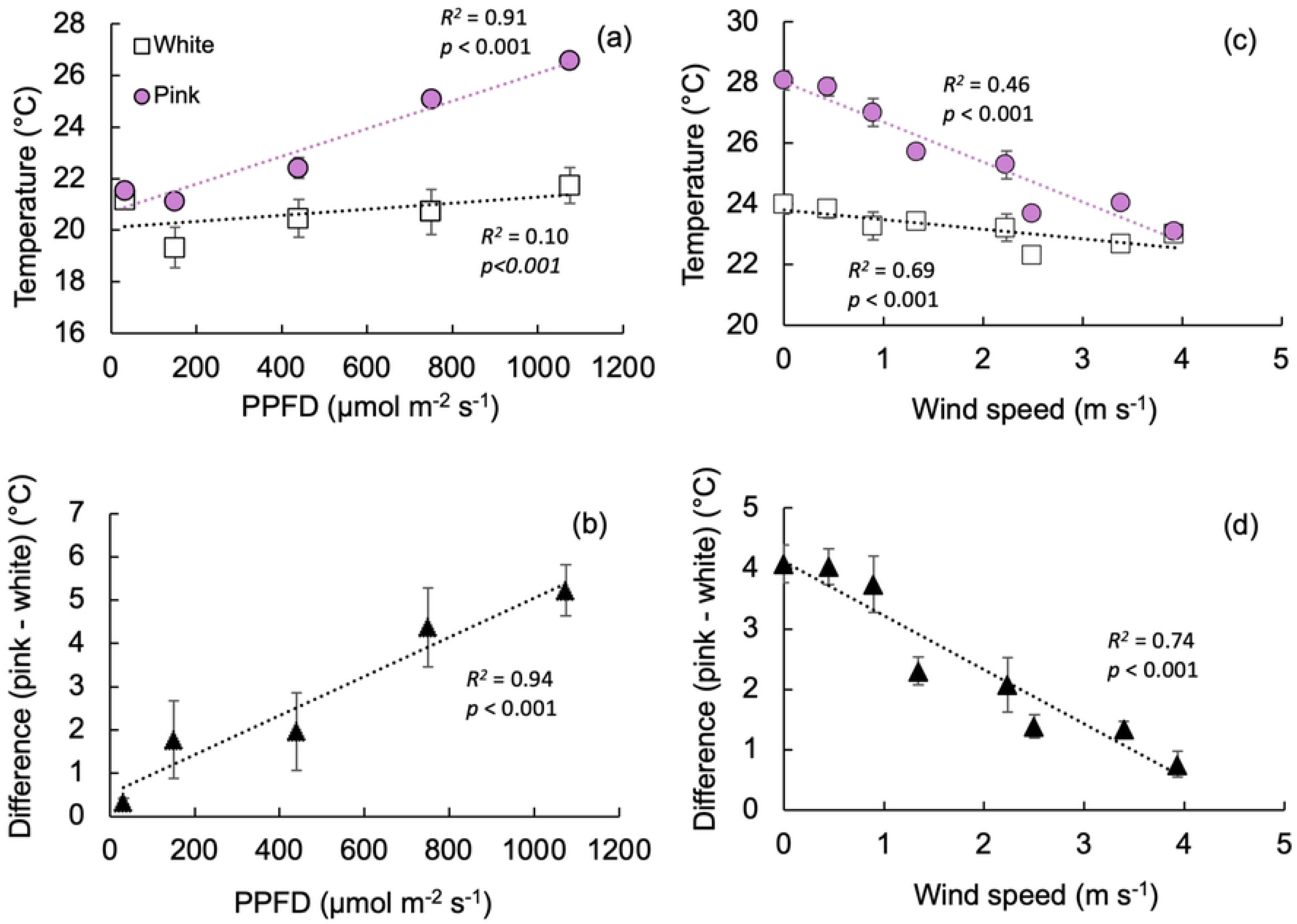
Effect of sunlight intensity and wind on difference in flower temperature between dark pink and white *Impatiens x hybrida.* Air temperature was 23 °C (± 1 °C), and wind was absent. Incident PPFD for wind experiment 1000-1100 μmol m^-2^ s^-1^. Points represent means of 5 replicates ± SE.

Wind had a significant cooling effect on both colors of flowers under high PPFDs (p<0.001 for both; Fig. 6c). Differences in flower temperature were greatest under low-wind speeds, and diminished gradually with increasing wind speed, until both color morphs reached an equilibrium with air temperatures at wind speeds of ∼4 m s^-1^ (Fig. 6d).

### In situ observations

Observations of intact plants with different colored flower cultivars occurring side by side consistently showed that darker color morphs were warmer than lighter colored morphs (Fig. 7). In many cases, white flowers were cooler than air temperature, while darker flowers were warmer. Differences in temperatures ranged from 7.4°C in pink versus white Sunpatiens*®,* up to 12.8°C in dark purple versus white cultivars of *Petunia x hybrida.* Darkly pigmented flowers were typically warmer than their leaves, while white flowers were either cooler than or similar in temperature as their associated leaves.

**Figure 7.**
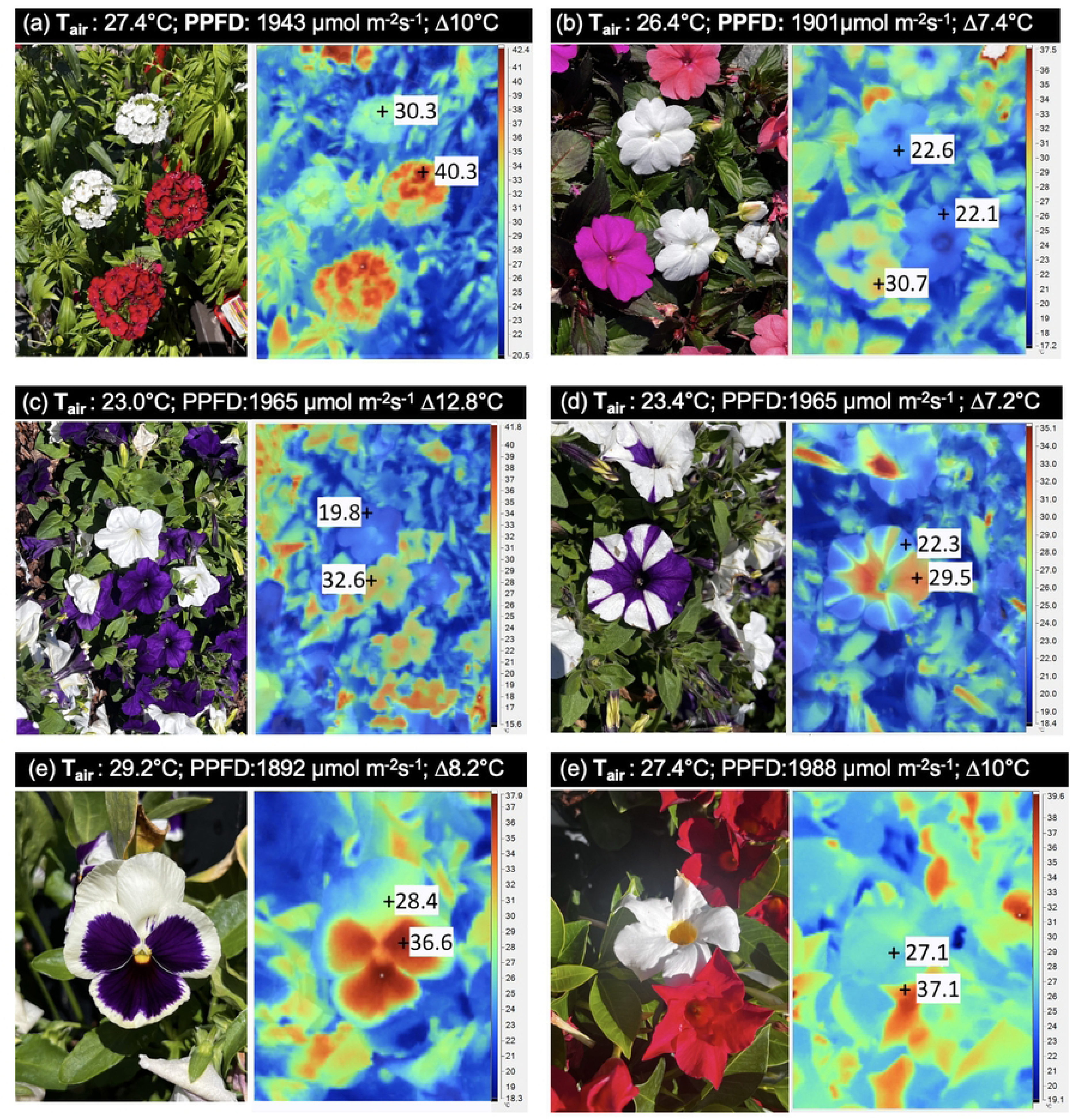
Field IR photographs of intact plants *in situ*. All images were taken under warm, clear sky, low wind (<1 m s^-1^) conditions during the summer of 2026 in High Point, North Carolina. Species shown include *Dianthus sp.* (a); SunPatiens*®, Impatiens x hybrida* (b); petunia, *Petunia x hybrida* (c, d); pansy, *Viola x wittrockiana* (e); rock trumpet, *Mandevilla sp.* (f).

## Discussion

The thermal ecology of flowers is important to understand because temperature directly impacts several important biochemical, physiological, and ecological factors associated with pollination, plant reproduction, and ultimately, seed/crop yield. In the context of climate change, identifying traits which confer tolerance to rising global temperatures is of utmost importance [66]. Yet, conflicting reports in the literature regarding the capacity for floral pigments to elevate flower temperatures have resulted in uncertainty surrounding this issue [41]. In the present study we used different color morphs of the same species to control for effects of morphology on flower temperatures and ran experiments under controlled environmental conditions to avoid major biophysical confounds. Our results clearly showed that flowers colored by red or purple anthocyanin pigments are consistently, and often substantially, warmer than white conspecifics. This trend was observed in 13 different plant species belonging to 13 different families, either tested using excised flowers in experiments or intact plants *in situ*.

*In situ* measurements further revealed that, while white flowers were commonly cooler than air temperature (likely due to evaporative cooling) and similar in temperature to leaves, darker colored flowers tended to be several degrees above air temperature. The average temperature difference between lightly-pigmented and white morphs used in controlled experiments was 2.9°C, while more darkly-pigmented morphs averaged 5.3°C (Table 1). While mean temperature differences are consistent with those cited for other species in the literature (see *Introduction*), many of our values were much higher. In controlled experiments, darker morphs of *Catharanthus roseus*, *Impatiens x hybrida* SunPatiens®. *Viola x wittrockiana* were roughly 8°C warmer than white morphs, and multiple taxa observed *in situ* exhibited temperature differences exceeding 10°C. Accuracy of temperature measurements made using the infrared camera were compared to temperatures using fine-wire thermocouples, and values were virtually identical (r^2^=0.996; Supplementary Fig. 1).

In addition to warmer petals, darker-colored flowers also had warmer central reproductive tissues than white flowers. As mentioned previously, multiple reproductive processes are sensitive to temperature, including flower opening, vaporization of pollinator-attracting volatiles, pollen development and germination, pollen tube formation, ovule development, and fertilization [30, 31, 32, 33, 34, 35, 36]. Hence, flower color could be a vital factor linked to success of native and agricultural species, especially those growing in climates close to their thermal maxima.

Because most cultivars tested did not significantly differ in temperature under UV inclusion/exclusion, we attribute warming effects primarily to absorbance of visible light. However, UV treatment did have a significant impact on temperature in the overall model, and some species were up to 2°C warmer under UV inclusion than exclusion. Such differences could be due in part to the presence of molecular side groups which increase molar absorptivity in visible and UV wavelengths of anthocyanins, as in [47]. However, some species showed differences in temperature under UV+/-in white flowers as well. Since all polyphenols absorb strongly in the ultraviolet wavelengths [45], these differences could be the result of warming by other (UV-absorbing) pigments besides anthocyanins.

### Interspecific differences

There are several possible explanations for enhanced warming in some species compared to others. In addition to higher concentrations of anthocyanins pigments, convex epidermal cells are known to increase absorptivity of photons by focusing light internally [67]. Kevan [40] reported that the enhanced floral warming in poppies could be linked to dense, prism-like ridges on petals composed primarily of cell wall. We observed raised, triangular dermal papillae in all three species with temperature differences at or exceeding 8°C, as well as in florets of *Pentas lanceolata* and *Salvia microphylla*. In addition to potentially focusing sunlight, dermal structures such as papillae and ridges can enhance or inhibit pollinator traction [68, 69], repel water [70], and promote warming by trapping heat through boundary layer effects [48]. Conversely, smooth and glossy petal surfaces have been shown to be more reflective than conical and silky appearing surfaces [71, 72]. Hence, petal surface structure interacts with and mitigates petal temperatures in complex ways.

Zooming out in scale, certain flower morphologies can promote warming by absorbing incoming sunlight and trapping the resulting longwave infrared radiation outside of wind flow within larger boundary layers associated with larger flowers, bowl or bell-shaped parianths, or closed structures resembling ‘micro-greenhouses’ [41, 48, 73, 74]. While there was no statistically significant effect of dermal structures or flower diameter on flower temperature in our larger model, this may be partly because of the low wind conditions utilized in our controlled study. Flowers in our controlled experiments were inserted to small holes within a white cardboard sheet, rendering many flush with the white surface. This effect would increase boundary layer resistance, as would the low-wind conditions under which the experiments were being conducted. Because all flowers were contained within these larger boundary layers, the influence of smaller boundary layer effects induced by dermal structures and flower diameter may have been dwarfed. Similarly, petal thickness did not have a significant effect on temperature in our model, even though thicker petals reduce reflectance by increasing internal scattering [75]. These factors illustrate the limitations of our controlled experiments, which were effective insofar as they isolated effects of petal color, but at the cost of minimizing the effects of other important morphological and biophysical factors.

The cooling effects of wind were demonstrated in an additional experiment using excised dark pink and white flowers of *Impatiens x hybrida* SunPatiens®. In the absence of wind, pigmented cultivars of were on average 4°C warmer than white morphs. Increasing wind speed reduced the temperature difference further, and the two reached the same temperature at wind speeds at or above 4 m s^-1^. Dietrich and Körner [76] similarly reported warming of alpine flowers only at wind speeds below 2-4 m s^-1^, and according to Grace [77] boundary layer resistance becomes negligible at wind speeds of 3-5 m s^-1^. These findings indicate that the most dramatic *in situ* warming effects driven by flower color would likely be observed in species in sunny climates with growth habits (or in habitats) that minimize wind flow, e.g., plants growing in dense stands or close to the ground.

### Ecological implications

In addition to impacts on reproduction, warmer petal temperatures also accelerate evaporation and transpirational water loss. While increased evaporative cooling may offset floral heating under drought stress as in [26, 78], stomatal closure will limit evaporative water loss, leading to excessive and potentially lethal temperatures, turgor loss, petal wilt, and ultimately negative impacts on floral display to pollinators. Although higher thermal tolerance has been reported in darker versus lighter color morphs of *Vinca minor,* warming effects may nevertheless exceed thermal limits, and/or narrow thermal safety margins [26]. It is also worth noting that greater transpiration and evaporation in darker flowers (attributed to warmer temperatures) likely offset some of the warming rendered by the pigments observed in this study, similarly to [26]. It is therefore likely that temperature differences associated with anthocyanin pigments could be even greater than those reported here.

Even if ample water is available, leaves and flowers of many species orient parallel to incoming irradiance, to minimize direct incident radiation and avoid overheating [48, 79]. We anecdotally observed a similar phenomenon in *Petunia x hybrida* flowers on a hot summer day, when bells of purple petunias were observed orienting parallel to incoming radiation, while white cultivars were oriented perpendicularly (Fig. 8). Such a dramatic change in flower orientation could impact pollinator visitation by making flowers less conspicuous, or by reducing spread of volatile attractants.

**Figure 8.**
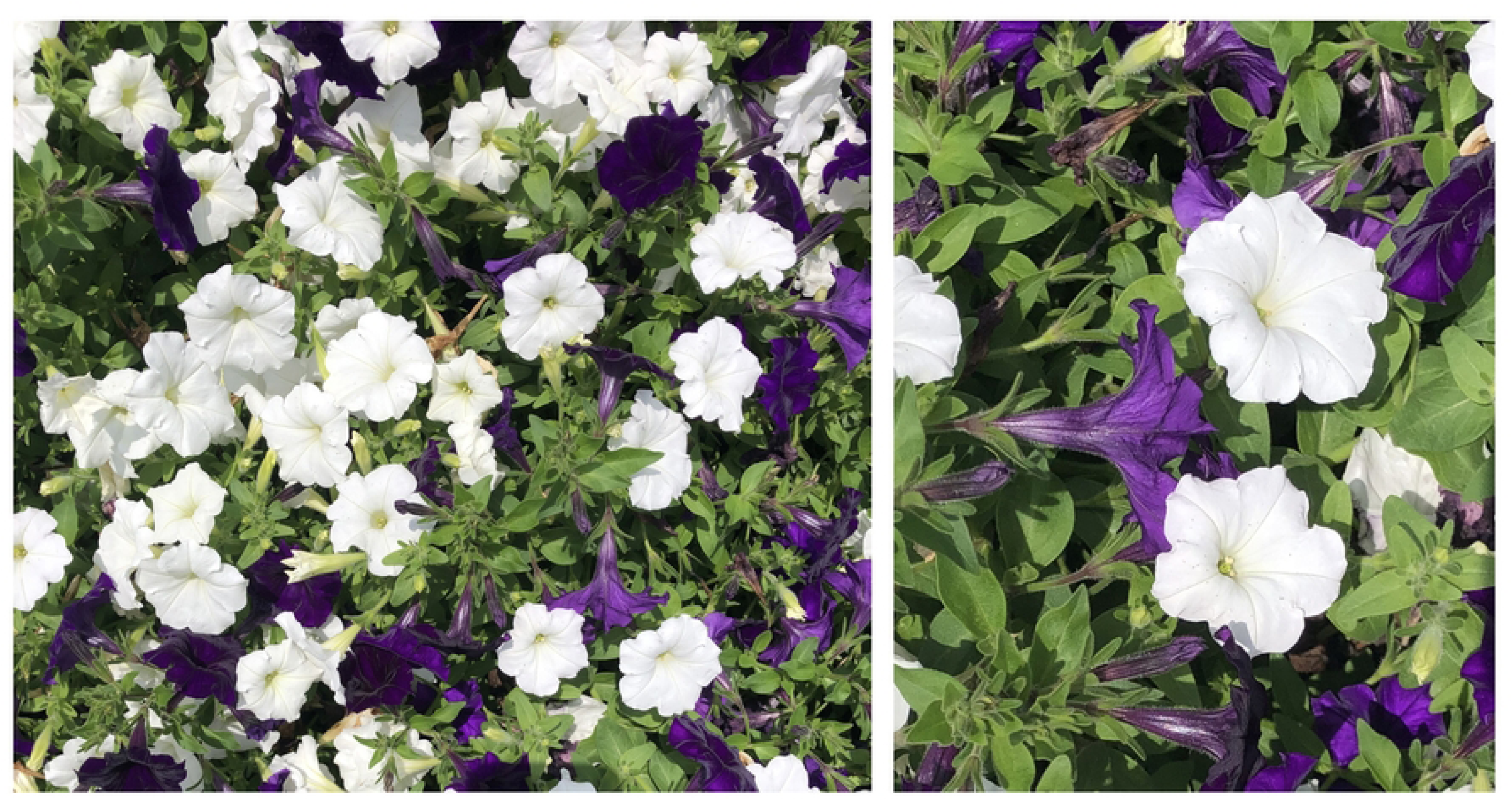
Differences in floral display angle observed *in situ.* Photograph was taken on 1 July 2023 at 13:15 during a cloud gap under partly cloudy conditions. PPFD was 1826 µmol m^-2^ s^-1^, air temperature was 35°C.

We also anecdotally observed that morning dew was more abundant on white flowers than darkly pigmented morphs, consistent with increased evaporation associated with warmer surfaces (N. Hughes personal observation). Evaporation of dew or raindrops from petals reduces vulnerability to fungal growth, burn spots, and blockage of stomatal gas exchange [80]. Accelerating the drying process in flowers would be especially beneficial in moist locations with high relative humidity, in which case elevating tissues above air temperature would help increase the leaf to air vapor pressure deficit and promote evaporation. No studies to our knowledge have tested the hypothesis that more darkly colored flowers dry more quickly than white flowers. Conversely, species which rely on direct foliar uptake of fog water to maintain optimal water balance, e.g., [81, 82], might be negatively affected by more rapid evaporation rates. Unfortunately, the vast majority of studies on such species focus on foliar uptake through leaves or fog drip, and no studies to our knowledge have tested whether plants can directly absorb fog water through flowers.

While in some cases floral warming may be an important selective pressure driving evolution of floral anthocyanins (e.g., in colder climates), in others it may be a side effect of some other function. Selection on pleiotropic traits associated with anthocyanin structural genes, such as number of flowers, seeds, defensive compounds, and biomass, could also play a role [15, 83, 84, 85]. This could potentially explain why colorful morphs of some species are more successful under arid conditions [86 and citations therein], even though increased floral pigments in dry environments seems counter-intuitive. It is unlikely that anthocyanins function as osmolytes in these systems, given their metabolic costliness relative to other solutes (e.g., simple sugars, ions), and their striking red color, which is unnecessary for an osmotic function [87].

### Flower color and climate change

The implications of our study for angiosperms in natural and agricultural systems are especially important in the context of climate change. In regions where plants and their pollinators already exist close to their thermal maxima, darkly-pigmented flowers may be at greater risk of overheating themselves and/or their pollinators. Examples of old and new world crops with red to purple flowers are numerous, and several examples are shown in Fig. 9. As previously mentioned, darker-colored flowers may also be more prone to desiccation stress due to increased evaporation, which may also lead to reproductive failure. These consequences may be offset to some degree by increased transpirational cooling [26, 78], as well as pleiotropic advantages associated with pigmented flowers under drought stress [86 and citations therein], although these latter mechanisms are not yet fully understood.

**Figure 9.**
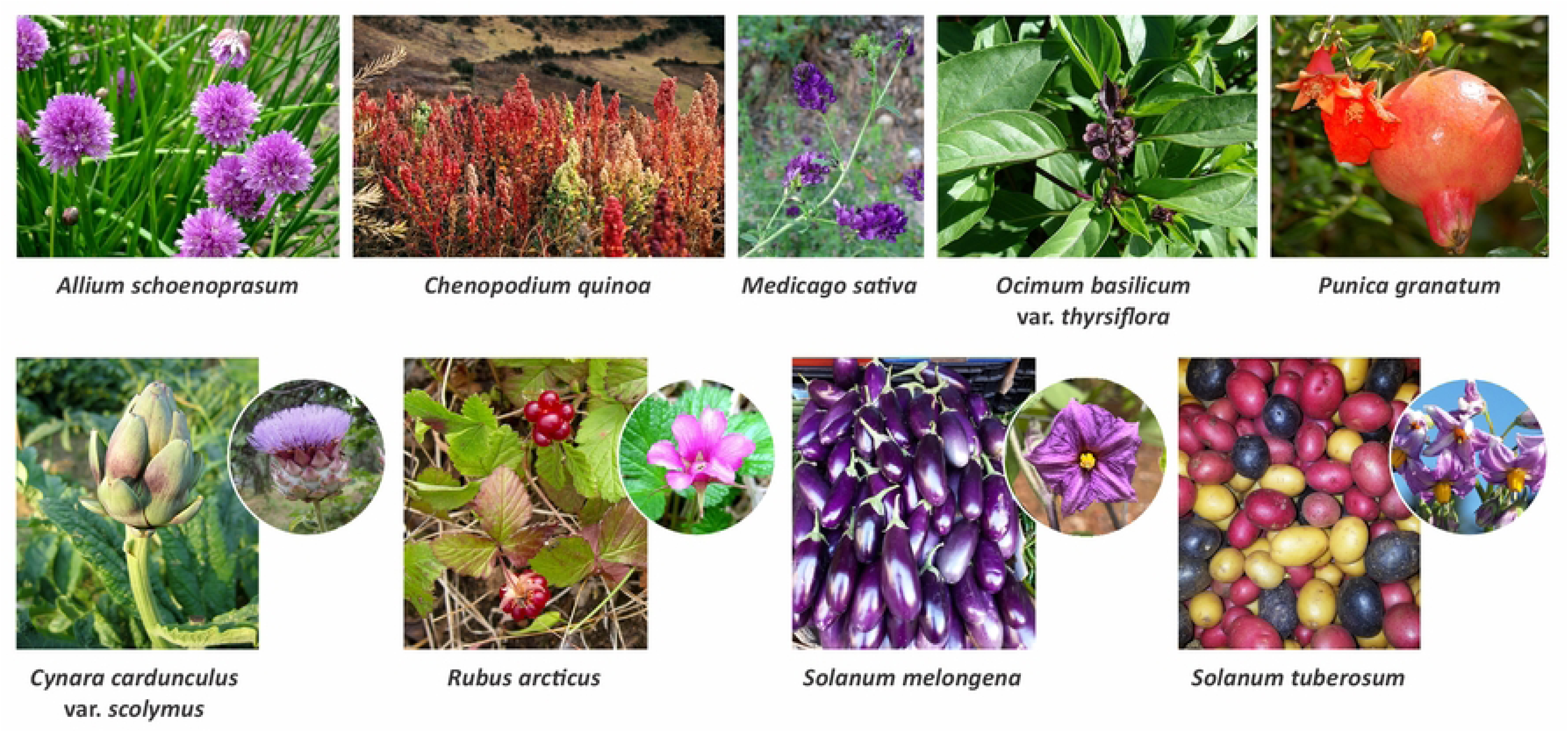
Examples of crops with red or purple flowers.

Rising temperatures associated with climate change have not been homogeneous across the globe. In the Arctic, temperatures are increasing at a faster rate (nearly 4x) compared to the rest of the planet [88]. In these latitudes, berries serve as an important summer food source for peoples and animals, and many of these species exhibit floral anthocyanins (e.g., *Rubus arcticus,* Fig. 8) [89, 90]. Warming effects in the Arctic may be offset somewhat during the day by increasing cloud cover [91, 92], which cool leaves and reduce evaporative water loss [93, 94]. In the tropics, subtropics, and midlatitudes, however, clouds have been decreasing [95, 96]. Decades of cloud models and observations indicate that stronger and deeper convection at the equator is narrowing the tropical rain belt, leading to fewer clouds and shorter rainy seasons in some parts of the tropics [95, 97, 98]. A stronger ascension at the equator is also a factor influencing the poleward expansion of the subtropical dry zones, resulting in a reduction of clouds at their poleward edges [95, 99, 100, 101]. In such regions, reductions in clouds could exacerbate heat stress from already rising global temperatures, and exacerbate thermal stress in species with colorful flowers.

One implication of our findings is that warmer global temperatures could potentially drive directional selection for smaller, and/or lighter-colored flowers. Such trends have already been reported by Sullivan & Koski [86], who used 124 years of herbarium records to demonstrate a significant correlation between temporal changes in flower color polymorphisms and air temperature among 12 North American species. Changes in flower color and/or size driven by rising temperatures could be in opposition to (or congruent with) directional selection by other forces such as pollinators, herbivores, and pathogens. If oppositional selection occurs, warmer global temperatures could help maintain floral color polymorphisms within populations [21]. A smaller petal area could also counteract warming effects of anthocyanins by promoting wind flow and convective cooling [48]. Indeed, some studies have shown that white flower morphs tend to be larger than pigmented conspecifics [50].

## Conclusions

Overall, our findings support the concepts that *i)* floral pigments can significantly increase flower temperatures, *ii)* these effects are substantial enough to potentially impact plant fitness, and *iii)* flowers with more pigmentation could be more vulnerable to overheating as temperatures increase under climate change. These findings are important because human agriculture, as well as the horticulture industry, depend on many plants with colorful flowers. Furthermore, changes in flower color and/or size can have community-scale effects, influencing pollinators, herbivores, and pathogens. For this reason, flower color should be considered an important trait linked with the potential success (or peril) of certain morphs, cultivars, or species, in a warming world.

## Acknowledgements

We thank Dr. Sandra Cooke for donating the Courtguard and Aclar, and High Point University curator of grounds Emma Martone for helping identify campus plants used in this study. Financial support to Elizabeth Ragan was provided by the High Point University Natural Science Fellows program. Photos in Figure 2 not taken by Nicole Hughes were from Wikimedia commons via Bojan Cvetanović (wax Begonia); KENPEI (baby sage shrub); Dave Whitinger (crepe myrtle panicle). Photos in Figure 9 from Wikimedia commons via Maurice Chédel (quinoa), Fructibus (eggplant), Daderot (potatoes), artichoke flower (cropped) (Stan Dalone & Miran Rijavec), Forest Service Alaska Region, USDA (arctic raspberry flower), Kristof Zyskowski & Yulia Bereshpolova (arctic raspberry plant), Donar Reiskoffer (chives), Forest & Kim Starr (Thai basil), Robert Flogaus-Faust (alfalfa). NMH planned and designed the research. SJF, EDR, and NMH performed experiments and conducted field work. NMW, JWC, SJF, and EDR analyzed data. JWC, EDR, SJF, NMH created figures. NMH and JWC wrote the manuscript. SJF, EDR, and NMW edited the final manuscript.

## Data Availability Statement

The data that support the findings of this study are openly available in Ag Data Commons. at https://doi.org/10.15482/USDA.ADC/27229290.v1.

## Supporting Information Captions

**Supplementary** Figure 1. Comparison of temperatures measured by the Fluke TiR110 thermal imaging camera with and fine-wire thermocouples connected to a S220-T8 eight channel datalogger (Huato, Shenzhen, China). Measurements were taken using beakers of water representing a range of temperatures between 0-40°C.

## Notes

### Competing Interest Statement

The authors have declared no competing interest.

